# AutoPhy: Automated phylogenetic identification of novel protein subfamilies

**DOI:** 10.1101/2022.07.21.501007

**Authors:** Adrian Ortiz-Velez, Jeet Sukumaran, Scott T. Kelley

**Affiliations:** Bioinformatics and Medical Informatics Program, San Diego State University, San Diego, CA, USA; Department of Biology, San Diego State University, San Diego, CA, USA

**Author notes:** Department of Biology 5500 Campanile Drive, Mail Code 4614, San Diego State University, San Diego, CA 92182, USA. Membership list can be found in the Acknowledgments section.

## Abstract

Phylogenetic analysis of protein sequences provides a powerful means of identifying novel protein functions and subfamilies, and for identifying and resolving annotation errors. However, automation of functional clustering based on phylogenetic trees has been challenging, and most of it is done manually. Clustering phylogenetic trees usually requires the delineation of tree-based thresholds (e.g., distances), leading to an *ad hoc* problem. We propose a new phylogenetic clustering approach that identifies clusters without using *ad hoc* distances or other pre-defined values. Our workflow combines uniform manifold approximation and projection (UMAP) with Gaussian mixture models as a k-means like procedure to automatically group sequences into clusters. We then apply a “second pass” clade identification algorithm to resolve non-monophyletic groups. We tested our approach with several well-curated protein families (outer membrane porins, acyltransferase, and dehydrogenases) and showed our automated methods recapitulated known subfamilies. We also applied our methods to a broad range of different protein families from multiple databases, including Pfam, PANTHER, and UNIPROT. Our results showed that AutoPhy rapidly generated monophyletic clusters (subfamilies) within phylogenetic trees evolving at very different rates both within and among phylogenies. The phylogenetic clusters generated by AutoPhy resolved misannotations, determined new protein functional groups, and detected novel viral strains.

## Introduction

The advent of deep sequence technologies, and the subsequent drop in the price of sequencing, has dramatically increased the rate of data acquisition for genomics, metagenomics, and transcriptomics. This development required the creation of faster and more accurate computational and bioinformatics algorithms, as well as a need for databases and search tools that can rapidly identify gene functions. Perhaps the most widely used method for protein function and family identification is the profile Hidden Markov Model (HMM), a Markov chain Monte Carlo sequence site transition model built from distantly related proteins of similar function with sequence gaps included in the scoring [1]. Profile HMM algorithms such as HMMER [1] implemented in Pfam [2] provide rapid and accurate protein function identification and database search tools. While Pfam is largely concerned with broad functional identification of proteins at the family level, there is also a tremendous diversity of functions within Pfam families and many differentiated functional roles are collapsed within HMM models. These protein “subfamilies” share the basic function of the family, but also contain many unique properties that confer a different substrate specificity or metabolic role of the protein.

The diverse bacterial family of porins illustrates this point. Outer membrane porins (OMPs) are proteins that form pores or channels in bacterial membranes that allow the passive diffusion of a variety of soluble molecules such as sugars [3]. Some OMPs are highly selective and only diffuse certain molecules (e.g., maltose by maltoporin, oligosaccharides by OmpG, long-chain fatty acids by FadL) while others, known as general diffusion porins, have no substrate specificity, though they may favor certain ions [4]. Although OMPs are distantly related sequences with similar basic structure and function, there are very distinct OMPs within that have discrete and even novel functions [5]. Search methods that identify broad functional categories, such as profile HMMs, detect fundamental properties of all OMPs but miss a lot of their functional subfamily diversity.

In combination with these broad classification approaches, phylogenetic methods provide a potentially powerful means of identifying important functional subfamily diversity within these larger classifications because they can be used to determine well-supported monophyletic clades of protein sequences within these families. In addition, phylogenetic branch length information can be used to identify highly divergent sequences (including large insertions and deletions) indicative of novel functional structures or motifs that may confer distinct functional properties. Phylogenetic methods have been, and are currently being, used to identify and classify protein functions. For example, the TreeFam and PANTHER databases use phylogenetics to identify protein subfamilies by partitioning the trees. While these databases are very useful resources, both of them rely on manual curation by a team of biologists and are not automated. This clearly becomes problematic as datasets continue to increase in size. Moreover, these databases are largely not concerned with subfamily functional identification, but rather higher level annotations [6, 7].

There have been attempts to partition phylogenetic trees to identify subfamilies without manual curation. Sjolander 1998 developed an automated method based on optimizing an encoded cost of a MSA of a protein family. However, this method was not strictly based on phylogenetic trees and has not been substantially tested. Another method known as Secator used neighbor-joining trees (BIONJ [8]) to assign dissimilarity values to nodes, allowing them to be collapsed into subfamilies at significant dissimilarity values [9]. While Secator is relatively fast, BIONJ is a relatively unsophisticated method and phylogenetic approaches have improved significantly in speed and complexity since it was developed (e.g., the Bayesian phylogenetic tool BEAST [10]).

More recently, two different groups developed partially automated phylogenetic subfamily detection methods to partition monophyletic clades in already constructed phylogenetic trees. A recent paper by Balaban et al. (2019) used graph theory to traverse the tree and partition it into putative subfamilies at the edges given a distance threshold (alpha parameter) [11]. Another method called PhyCLIP determines clusters in a phylogenetic tree using an optimization equation with parameters such as a minimum number of sequences to solve the partitioning problem [12]. Although these methods are technically automated, both require the selection of arbitrary parameters (like alpha and a minimum number of sequences) to group tips of the tree by partitioning at edges. Changing the parameters can drastically change results, requiring users to re-run the programs until they return ‘expected’ results, leading to a classic *ad hoc* problem.

Here, we present a new approach for automated phylogenetic identification of protein subfamilies called AutoPhy. AutoPhy works with any phylogenetic estimation method and substantially improves previous automated tree-based subfamily clustering approaches by circumventing the need for an *ad hoc* parameter selection during the estimation process. Our approach combines phylogenetic analysis, non-linear dimension reduction, expectation-maximization, and monophyly resolution to identify discrete phylogenetic clusters using evolutionary branch length information. In addition, after identifying putative new subfamilies, AutoPhy includes methods for detecting highly divergent subfamilies which may be indicative of significant evolutionary functional constraints.

We validated AutoPhy with 17 protein families with highly divergent evolutionary rates both among and within the families. We also applied AutoPhy to identify novel phylogenetic clusters in rapidly evolving viruses. Overall, our results show that AutoPhy rapidly and automatically identifies discrete monophyletic clusters across a wide range of differentially evolving protein families, allowing detection of novel protein subfamilies, correction of annotation errors, and identification of unique sequence motifs.

## Materials and methods

Fig 1 details the automated phylogenetic clustering workflow developed in this study, and we refer to the different parts of this workflow diagram where appropriate throughout the method section. Our clustering methods can accept three possible sources of input: unaligned sequences (Figure 1: “Sequences”), multiple sequence alignments (“Alignments”), and pre-generated phylogenetic trees such as the NextStrain phylogenetic trees (“Phylogeny”).

**Fig 1.**
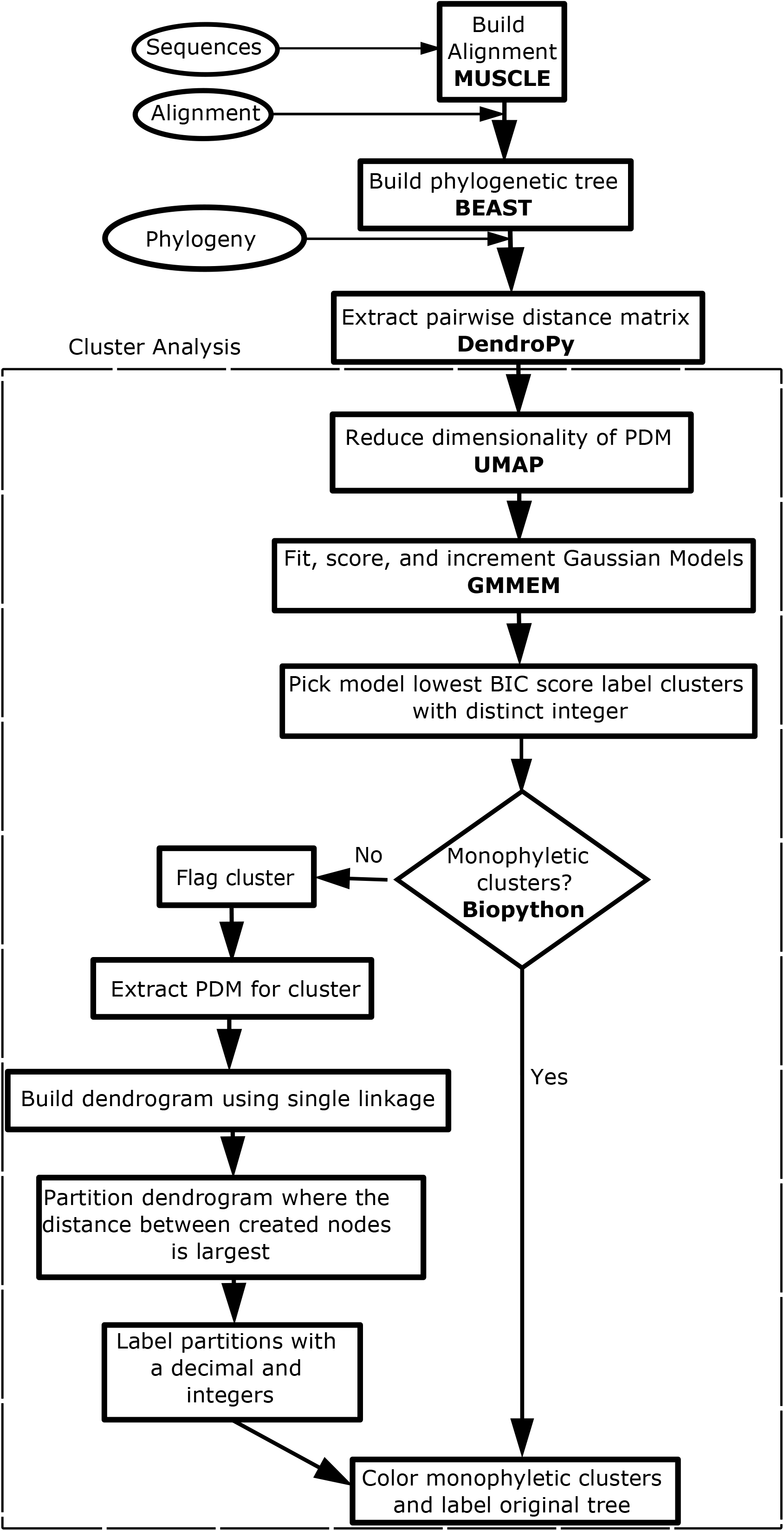
Flowchart : Flowchart of the clustering and feature analysis workflow. The workflow allows input of raw protein sequences from a fasta file, a protein MSA from a nexus/fasta file, or a fully constructed phylogenetic tree with branch lengths from a nexus/newick file. The analysis workflow was tested with raw sequences obtained from the PANTHER [7], UNIPROT [13], and KEGG [14–16] databases, MSAs from the Pfam [2] database and prebuilt phylogenetic trees come from Nextstrain [17] database. To speed up and reduce noise, UMAP is used to reduce the dimensionality of the extracted PDM from the natural tree lengths. To identify putative clusters, a Gaussian Mixture Model was built to find the best fit combination of Gaussian peaks on the PDM with reduced dimensionality. A second pass is added to partition Gaussian peaks from the best fit model that do not relate to a clade on the tree, to maximize the monophyletic clusters.

### Test datasets

Four protein family datasets obtained from Uniprot [14] were used as “positive controls” to validate the results our workflow because of their well-characterized functions and annotations: (1) A dataset of outer membrane porins (OMPs) obtained from UniProt (search term: “OMP”) that includes well-characterized porin proteins with different transport roles and unique motifs (e.g., OMPF, OMPC); (2) A dataset of acyltransferases obtained from UniProt (search terms: ‘steroid receptor AND reviewed:yes AND organism:”Homo sapiens (Human) [9606]”‘) that includes annotation information on specialized substrates and chemistry, (3) A dataset of dehydrogenases obtained from UniProt (search terms: ‘steroid receptor AND reviewed:yes AND organism:”Homo sapiens (Human) [9606]”‘) that includes finer annotations defining protein specific substrates and products, and (4) A dataset of human steroid receptor proteins obtained from UniProt (search term: ‘steroid receptor AND reviewed:yes AND organism:”Homo sapiens (Human) [9606]”‘) that includes protein specific ligand-binding information. In addition to the four “positive control” Uniprot datasets, the sequences obtained from the PANTHER [7] database had been manually curated and could also be used for validation.

After validation, we applied our workflow to analyze a variety of protein families representing a wide variety of enzymatic functions. These datasets included sets of bacterial, eukaryotic, and viral proteins with highly differentiated rates of evolution among the sets, as well as between clusters within datasets. Table 1 provides detailed information on all the protein families analyzed in this study, including the database where the sequences were obtained, the number of sequences and MSA positions and the accession numbers or search terms used to obtain the sequences. Sequences were obtained from 4 different databases: Uniprot [14], KEGG [15–17], PANTHER [7], and PFAM [2]. We also tested our clustering approach on 3 resolved viral phylogenetic trees from Nextstrain [REF]: Mumps, and Sars-CoV-2 (collected: Feb. 2022). The complete collection of datasets, including sequences, MSAs, and phylogenetic trees, as well as the diagnostic plots, analysis results, and programming code, can be obtained at https://github.com/aortizsax/autophy.

**Table 1.**
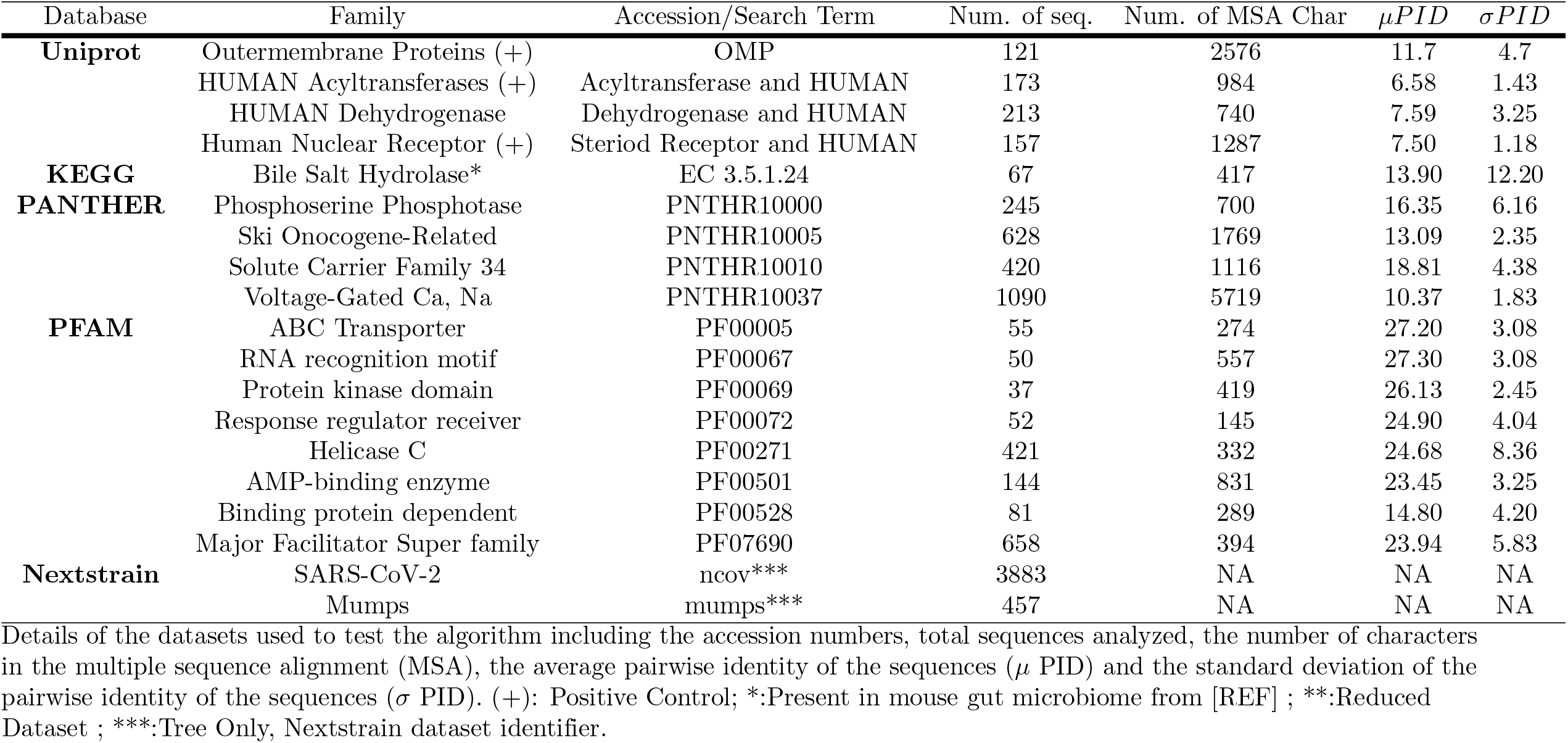
Details and statistics of the datasets used to test AutoPhy.

### Sequence Alignment and Phylogenetic Analysis

Multiple sequence alignments were performed using MUSCLE v3.8.31 [18] (Figure 1: “Build Alignment MUSCLE”). Phylogenetic trees of the data sets were inferred under a Bayesian framework using BEAST2 v2.6.3 [10] a Bayesian sampler Monte Carlo Markov Chain tree construction using the JTT site-specific model of protein evolution [19] and (gamma) category count of 6 to model 6 different site rates (“Build Phylogenetic Tree BEAST”) for a JTT+G site model. The Yule model was used to model topology and branch lengths in the analysis. This model assumes an unknown constant birth rate for every branch. Distributions of the priors used for the gamma shape distribution included uniform and exponential. For the viral datasets that did not have pre-generated phylogenetic trees (SARS-CoV-2 and Mumps; Table 1), we used an invariant site model to model the positions which do not change amongst sequences with a beta distribution prior with a mean of 0.9 for a JTT+I site model. The viral datasets did not require a gamma model to represent multiple site rates because the percent of invariant sites was so large that it coud be assumed there are only two rates of site changes those that do (uniform rate) and those that do not (no rate). The JTT+I model utilized for viral datasets was better than JTT+GI and JTT+G because JTT+I trace converges and mixes well as opposed to the others trace which did not mix well. The length of the Monte Carlo Markov Chain was adjusted based on the size of the dataset. First, the length of the MCMC was 10 million and sampled every 1000. If the trace did not converge or mix enough due to size of data set or complexity of model, the length of the MCMC was increased to 1 billion and sampled every 10,000 [20]. The first 10 % of the samples were discarded as burn-in following visual inspection in Tracer v1.7.1. [21] The program Tree Annotator v2.6.4 [22] was used to identify the maximum clade credibility (MCC) tree and to summarize the posterior of trees sampled by BEAST2 (Median node heights). Trees were viewed using FigTree v1.4.4 [23] and colored using the ETEtoolkit [24], Biopython [25], and TreeViewer v1.2.2 [26].

### UMAP dimension reduction and GMMEM Clustering

The first step in our automated phylogenetic cluster identification was the creation of a pairwise distance matrix (PDM). This matrix was calculated by parsing the branch length estimates of a given phylogenetic tree using the Python phylogenetic computer library DendroPy 4.5.1 [27] and summing the total distances between all pairwise leaves (Figure 1: “Extract pairwise distance matrix DendroPy”). Next, we used Uniform Manifold Approximation and Projection (UMAP) to reduce the dimensionality of the PDM to 4 dimensions for clustering (“Reduce dimensionality of PDM”). UMAP maintains the topology relationships between the rows in a data matrix while reducing the number of columns/dimensions to a size that humans can visualize (e.g., 2 to 3 dimensions), and to a size that eliminates noise for computers to cluster (3 to 10 dimensions). We then applied Gaussian mixture models fitted with Expectation maximization (GMMEM) to model Gaussian distributions where each distribution is a cluster of the reduced matrix (“Fit, score, and increment Gaussian Models GMMEM”). To determine the best fit number of Gaussian distributions or clusters are required to fit the data, models of increasing number of Gaussian distributions were scored using Bayesian Information Criterion (BIC). BIC is a model selection score that balances a model’s fit vs. its complexity (number of dimensions), with lower BIC scores optimizing this balance. The model with the lowest BIC score was selected and the clusters were labeled with a unique integer (“Pick model lowest BIC score label clusters with distinct integer”).

### Monophyletic Cluster Maximization

In some cases, the GMMEM model clusters were not monophyletic. The “monophyletic” function in Biopython v1.78 [25] was used to check whether all members of a particular cluster were part of the same monophyletic group (Figure 1: “Monophyletic clusters? Biopython”) and non-monphyletic clusters were noted (“Flag cluster”). In the case of a paraphyletic or polyphyletic cluster, we recursively extracted the largest monophyletic group or tree split using edge.bipartition.leafset as newick string() from DendroPy Phylogenetic Computing Library 4.5.1 [27] until all monophyletic groups in the polyphyletic groups had been identified. Once the cluster was split into monophyletic groups, unique decimal values were added to the end of the integer label, creating a floating-point label for all clusters that were previously non-monophyletic (“Label partitions with a decimal and integers”). At the end of this process, all the monophyletic clusters were colored for easy visualization (“Color monophyletic clusters”).

### Log ratio to evaluate evolution of subfamilies

To quantify the relative degree of intra-subfamily divergence, we develop a log ratio score that compared the within subfamily divergence to its subtending branch using Eq (1).

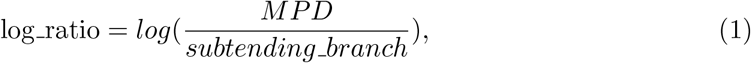

where MPD is the mean pairwise distance of the tips included in a cluster defined as Eq (2)

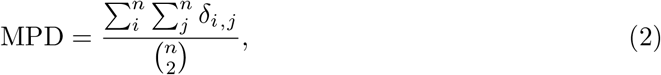

where i j, delta (i,j) is the phylogenetic distance between species i and j, and n is the number of species in the sample. We used DendroPy Phylogenetic Computing Library 4.5.1 [27] to calculate the MPD for putative subfamilies and parsing their subtending branches.

## Results

The pairwise sequence identity within the 23 protein families we analyzed in this study ranged from 6.58±1.43 to 99.03±2.31(percent) (Table 1). We first tested our approach using our “positive control” protein families: bacterial outer membrane porins (OMPs), bile-salt hydrolases (BSHs), human acyltransferases (AT), and human nuclear receptors (NR). First, to clarify our methodology, we provide detailed results for the automated phylogenetic clustering approach for the OMPs (Figs. 2-6), which is the same process used for all the protein families. Next, we summarize the rest of the protein family results. All the results described in this section are available along with the data and the programming scripts on github (https://github.com/aortizsax/clustree).

**Fig 2.**
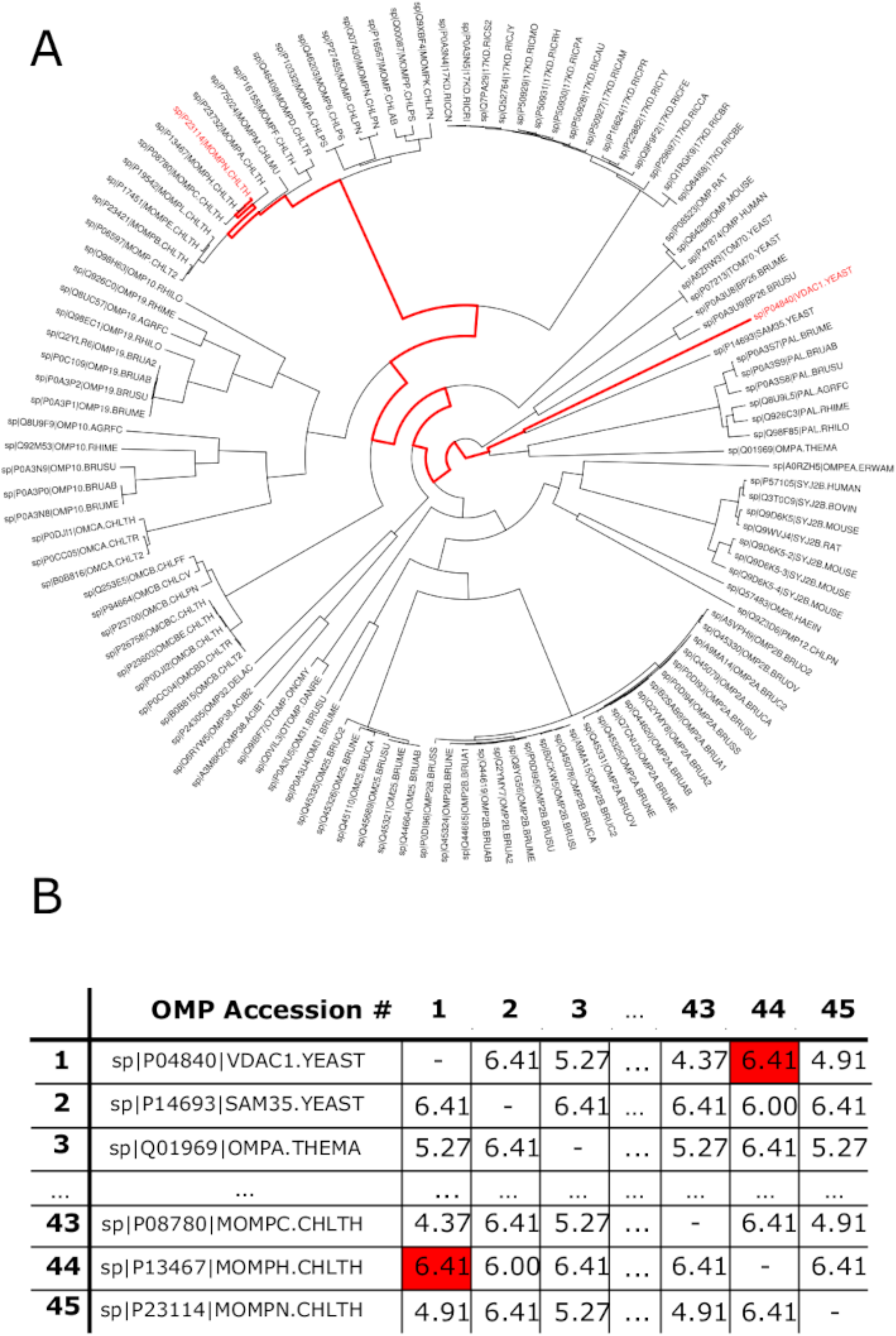
Pairwise Distance Matrix: Generation of PDM given a phylogenetic tree. (A) The phylogenetic tree of the outer membrane protein (OMP) sequences. (B) The PDM for the OMP based on the branch lengths of the phylogenetic tree. The numbers in the table indicate total pairwise branch length between along the phylogenetic tree each pair of proteins. The path from TOLC ECOLI to PMP10 CHLPN is highlighted in red in both A and B.

### Outer Membrane Porin subfamily identification

Bacterial OMPs form pores or channels in bacterial membranes that allow the passive diffusion of solutes such as sucrose [3]. Some porins are highly selective and only allow the diffusion of certain molecules (e.g., maltose by maltoporin, oligosaccharides by OmpG, long chain fatty acids by FadL) while others are known as general porins have no substrate specificity, though they may favor cations or anions [3]. The OMP dataset we tested was comprised of 121 sequences obtained from Uniprot (Table 1). Figure 2A shows the phylogenetic tree of these sequences obtained by BEAST as well as a portion of the PDM. The results of the UMAP analysis PDM is shown in figure 3A, and the results of the GMMEM analysis, which had a lowest BIC score of 16, is shown in figure 3B. The 10 peaks identified by the lowest BIC score from the GMMEM analysis formed clearly separable clusters on the UMAP graph (Fig. 4A). When relating those peaks back to clades on the initial tree, 4 out of the 10 identified clusters were monophyletic, while the remaining 6 were para- or polyphyletic groups (Fig. 4B). MCIM was used to resolve the remaining non-monophyletic groups (Fig. 5), which were divided into multiple different monophyletic groups (Fig. 6).

**Fig 3.**
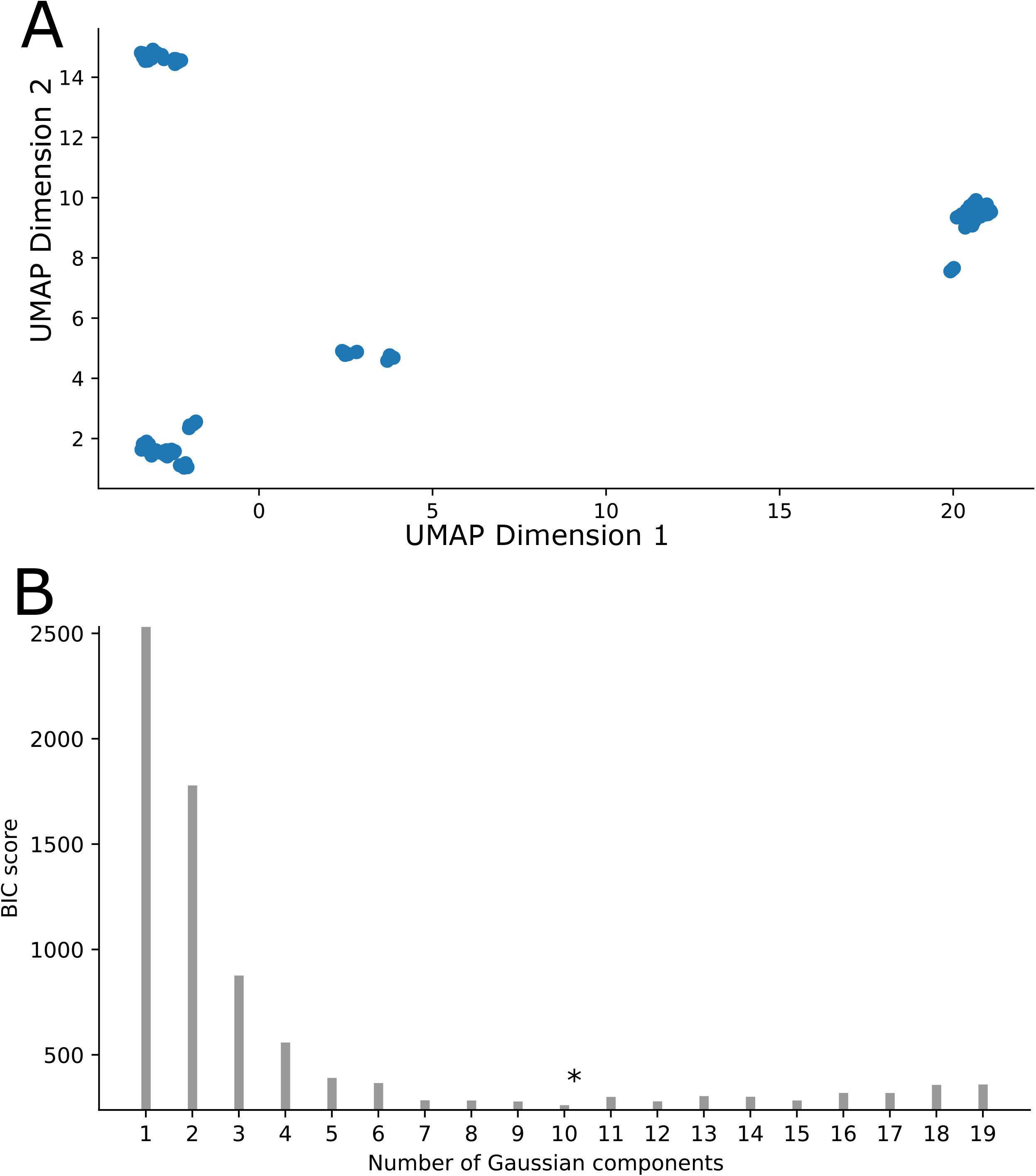
UMAP-GMMEM analysis of phylogenetic tree data: (A) UMAP visualization of unlabeled tips for the first 2 dimensions of the PDM from figure 1. (B) Scores for the GMMEM models for 1 to 19 clusters or peaks. In this example, the model with 10 clusters had the lowest BIC score.

**Fig 4.**
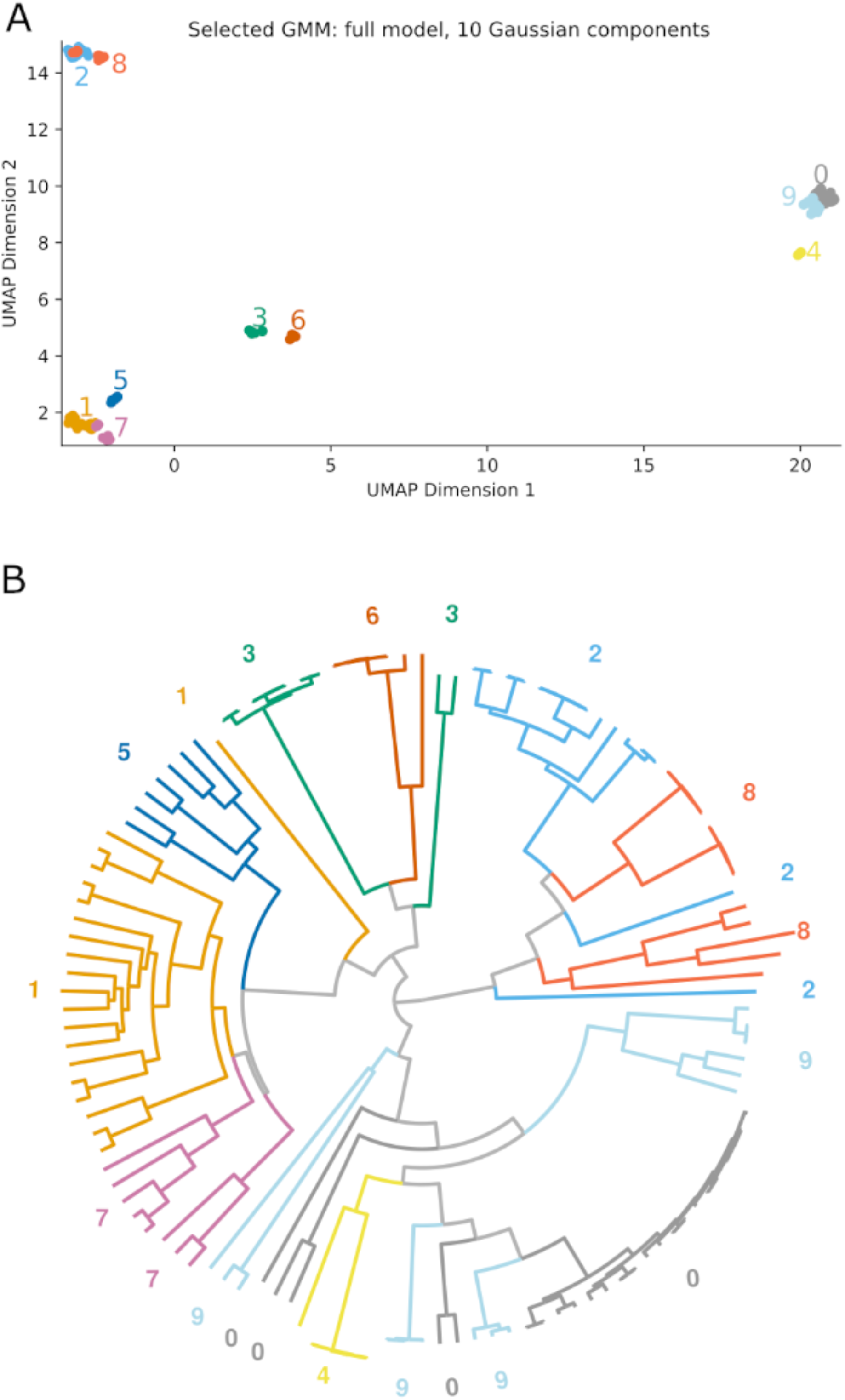
GMMEM Lowest BIC: Visualization of the lowest scored model identified by the GMMEM indicated by color on the (A) UMAP plot showing the best fit model’s 16 Gaussian peaks (B) Phylogenetic tree colored and labeled for their Gaussian peaks from the best fit model. The numbers correspond to each of the Gaussian peaks in the GMMEM model relating to the tree. Note that two clusters 0, 1, 2, 3, 7, 8, and 9 are polyphyletic and there is also a deep branching separation in cluster in 10.

**Fig 5.**
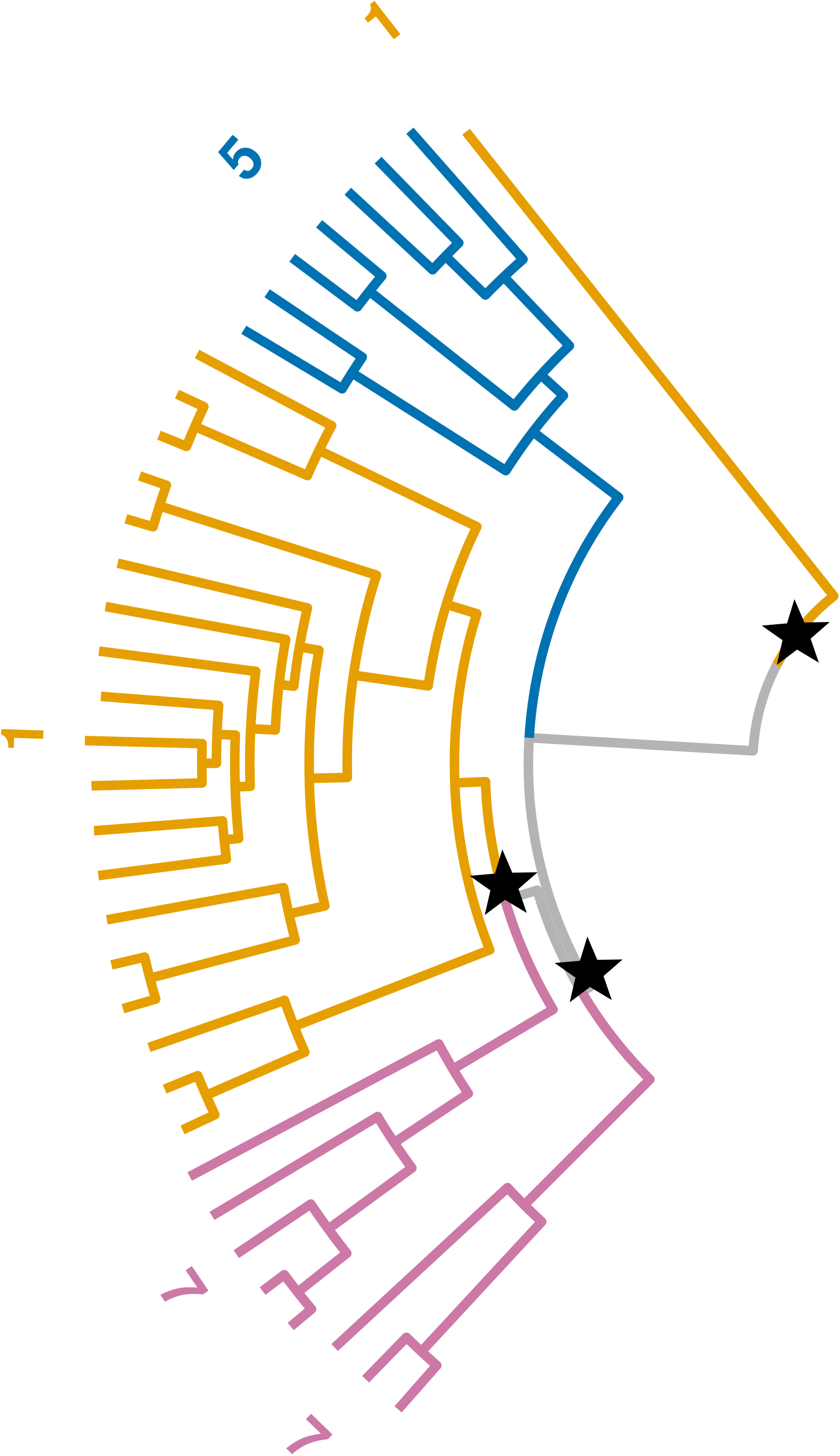
Monophyletic Cluster Maximization : MCIM on para-(/poly)-phyletic clusters. A subtree of OMP is shown performing MCIM on clusters 1 and 7 to resolve clusters from UMAP-GMMEM that do not form clades on the tree due to certain topology and branch length conditions. This decreases the minimum number of sequences for a group as well as leads to all sub-clusters corresponding to clades/monophyletic groups.

**Fig 6.**
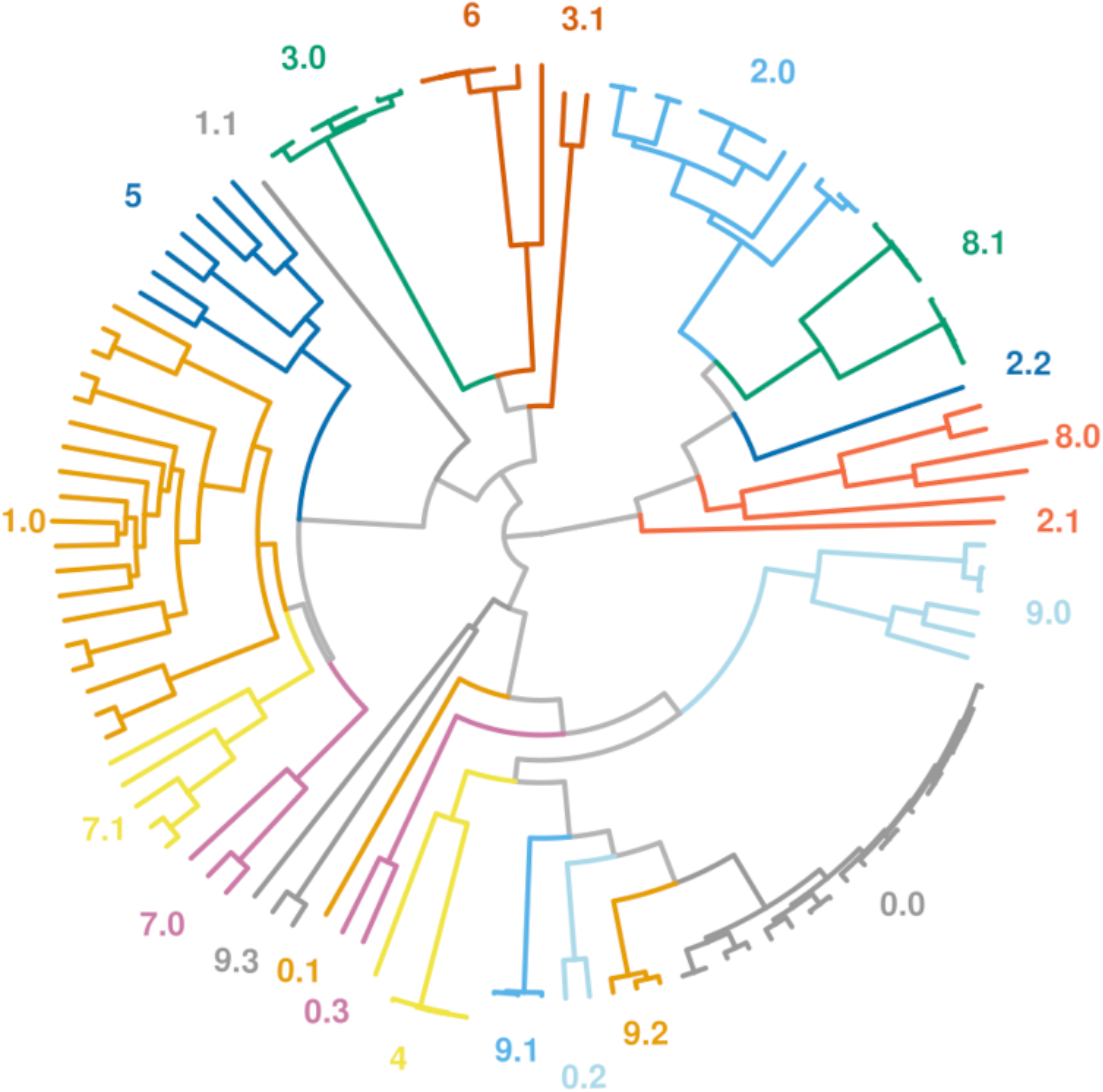
Final Result of UMAP-Gaussian Mixture Model-Monophyletic Cluster Maximization: Full algorithm UMAP-GMMEM-MCIM results for the OMP phylogeny are shown by color and number. Labels with an integer are clades that clustered with UMAP-GMMEM. Labels with a floating-point number are clades that needed refinement via clade identification of the resolved para(poly)-phyletic UMAP-GMMEM cluster.

### Bile salt hydrolases

The bile salt hydrolases are a family of proteins that deconjugates glyco-conjugated and tauro-conjugated bile acids and maintains in intracellular homeostasis and modifications in the architecture and composition of the cell membrane. Several studies indicate that BSH activity affects both the host physiology and the microbiota; [8] suggested that BSH could play an important role in the colonization and survival of bacteria in the gut. Also, recent work has shown that BSH and free BA participate in a variety of metabolic processes that include regulation of dietary lipid absorption, cholesterol metabolism, and energy and inflammation homeostasis. The sampling set used comes from a mouse model of PCOS [29]. We used our method to analyze 67 sequences belonging to BSH proteins downloaded from KEGG (Table 1). Figure 9A shows the initial tree obtained by BEAST with the GMMEM peaks indicated (Low BIC Model=9 peaks). The UMAP-GMMEM clusters numbered 1 and 8 proved to be non-monophyletic and after MCIM, each of those GMMEM clusters were split into 2 new monophyletic groups, (Fig. 7B).

**Fig 7.**
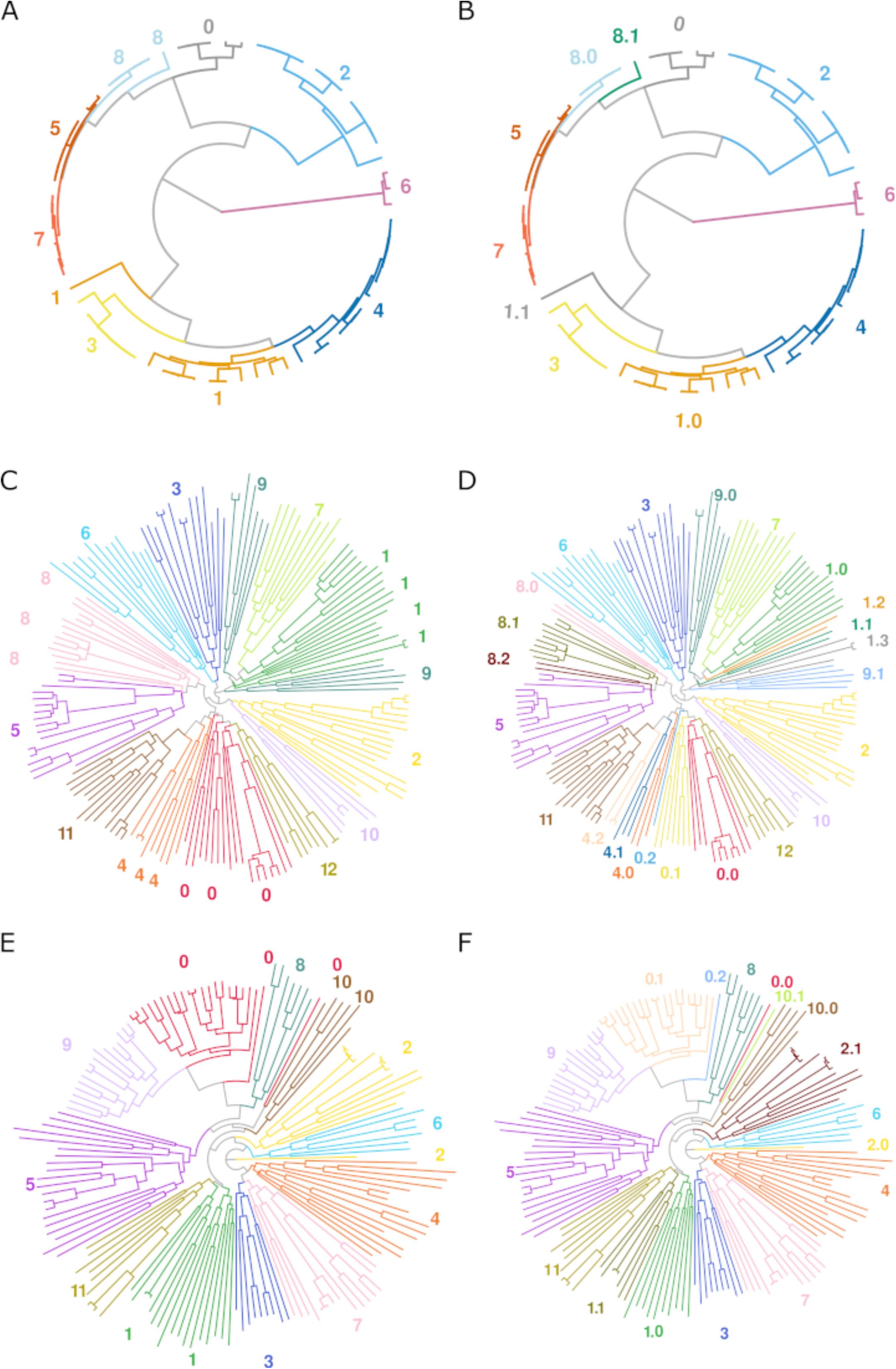
Post GMMEM and Final clustering results: BSH family(A,B), AT family(C,D), and NR family(E,F). Full algorithm UMAP-GMMEM-MCIM results are shown by color and number. Labels with an integer are clades that clustered with UMAP-GMMEM(both columns). Labels with a floating-point number are clades that needed refinement via clade identification of the resolved para(poly)-phyletic UMAP-GMMEM cluster(right column).

### Acyltransferase

The acyltransferase proteins represent an important protein family which transfers acyl groups (group of ketone with a chain of carbons) to other molecules involved in phospholipid biosynthesis [30]. For example, circulation of cholesterol away from tissues is dependent on the activity of lecithin cholesterol acyltransferase (LCAT). LCAT is a water-soluble enzyme that catalyzes cholesterol and phosphatidylcholines (lecithins) to cholesteryl esters and lyso-phosphatidylcholines on the surface of hydrophobic high-density lipoproteins(HDL) [31]. Figure 9C shows the initial tree obtained by BEAST of 173 acyltransferase protein sequences obtained from Uniprot (Table 1). After UMAP-GMMEM analysis, we identified 4 different clusters (Low BIC Model=13 peaks), although only 5 of these proved to be monophyletic. After MCIM, there were a total of 23 monophyletic clusters (Fig. 9D).

### Nuclear Receptor

The Human Nuclear Receptor proteins are transcription factors that are ligand inducible by small lipophilic molecules such as steroid hormones, retinoids, and free fatty acids and control many of the reproductive, developmental, and metabolic processes in eukaryotes. The mediators of these effects are nuclear receptor proteins, ligand-activated transcription factors capable of regulating the expression of complex gene networks [32]. Figure 7E shows the initial tree obtained by BEAST of 157 human nuclear receptors protein sequences obtained from Uniprot (Table 1). After UMAP-GMMEM analysis, we identified 12 different clusters (Low BIC Model=12 peaks), while 8 of these proved to be monophyletic. After MCIM, there were a total of 17 monophyletic clusters (Fig. 7F).

### Log ratio analysis

The log ratio of mean pairwise distance of a cluster by its subtending branch was obtained for all identified clusters before and after MCIM splitting. Our ratio was comprised of the MPD in the identified clusters and subtending branch to that cluster so a highly negative score would mean the MPD is very small compared to the its subtending branch. These negative log ratios for clusters means the cluster might have distinct in the evolutionary forces acting on those clusters. Figure 8a shows that before MCIM split the para/poly-phyletic clusters, the OMP were grouped together near zero. Figure 8b shows that after MCIM splitting, the clusters in the top (group 8 and 9) partitioned into clusters that shift to the middle of the graph or in some cases to the bottom where confident clusters are located. Figure 8c and 8d show the spread of the final clusters’ log ratio results of BSH and AT, respectively. Supplementary figures include the other highlighted subfamilies.

**Fig 8.**
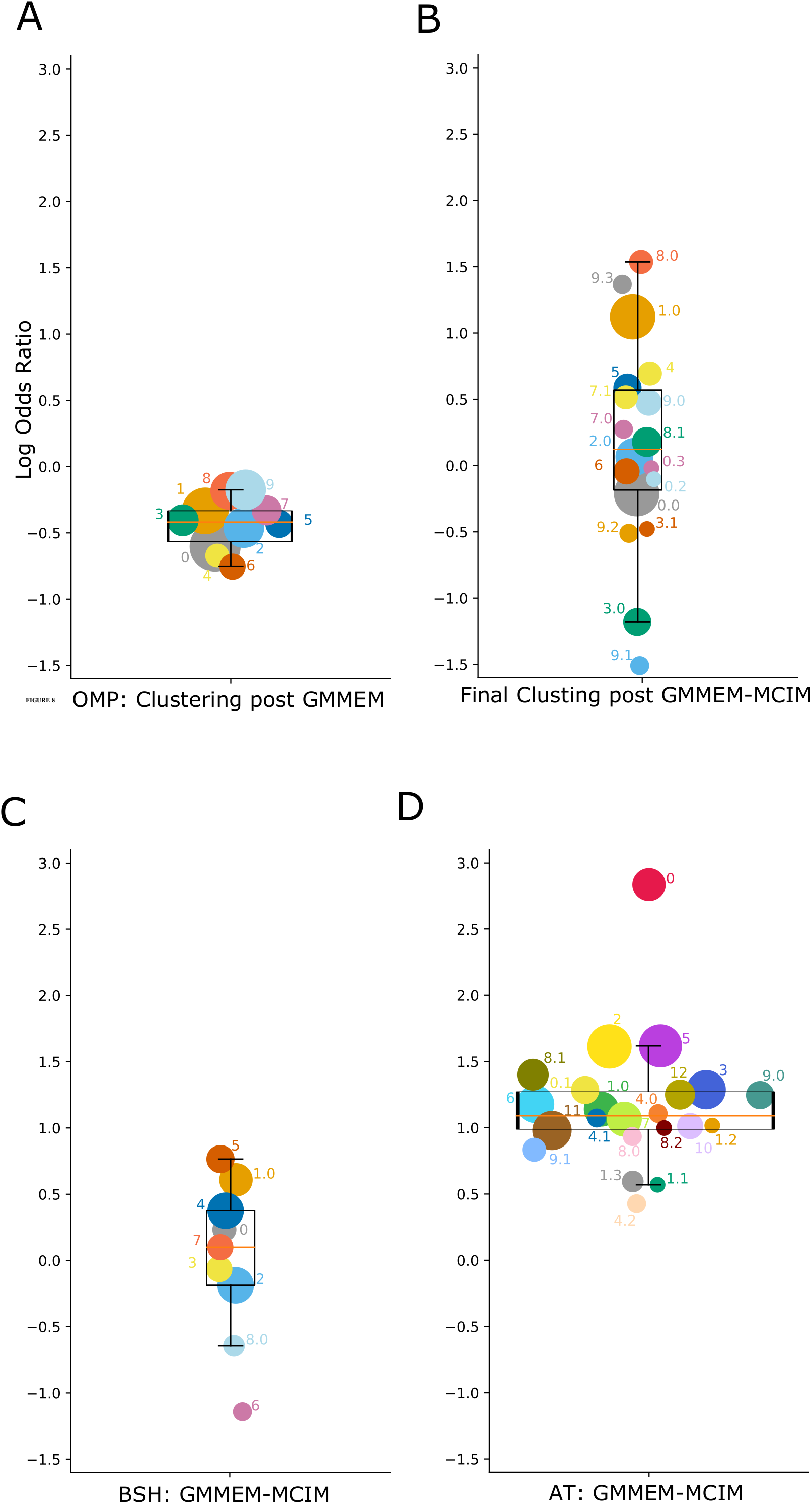
Log ratio analysis: Result of the log ratio analysis of phylogenetic clusters for different test data sets. The values represent the log of the ratio of the mean pairwise distance (MPD) within each phylogenetically determined subfamily to its subtending branch. Each subfamily log ratio is colored, label, and sized by its final clustering result. Log ratio values for post UMAP-GMMEM OMP(A); and the final clustering of OMP(B), BSH(C), and AT(D), respectively.

### Other protein families

Finally, we applied our pipeline to a broad range of different protein families, evolving at drastically different rates from a multitude of databases, including Pfam, PANTHER, and UNIPROT. Additionally to assess robustness against ultrametrtic trees, we tested our algorithm on two viral trees mumps and ncov with different phylodynamics. Table 2 summarizes the clustering and site analysis results on broad range of data sets. The table presents the number of putative subfamilies and the time needed to cluster the original phylogenetic tree. The number of putative subfamilies is synonymous with the number of clusters. The time to cluster the original tree is comprised of the time needed from PDM creation to coloring the final tree. The last column of table 2 show the percent of independent sites by Fisher exact test on subset of the top 10% divergent sites.

**Table 2.** Summary results for 21 datasets analyzed used to test AutoPhy.

| Database | Family | Num. of putative subfamily | Time to cluster tree (sec) |
| --- | --- | --- | --- |
| Uniprot | Outermembrane Proteins (+) | 22 | 4.77 |
|  | HUMAN Acyltransferases (+) | 23 | 7.78 |
|  | HUMAN Dehydrogenase | 26 | 7.71 |
|  | Human Nuclear Receptor (+) | 17 | 7.17 |
|  | Ebola S Glycoprotein | 19 | 6.60 |
|  | SARS-CoV2 S Glycoprotein | 51 | 22.00 |
| KEGG | Bile Salt Hydrolase (+) | 11 | 4.68 |
| PANTHER | Phosphoserine Phosphatase (PNTHR10000) | 10 | 5.37 |
|  | Ski Onocogene-Related (PNTHR10005) | 23 | 3.88 |
|  | Solute Carrier Family 34 (PNTHR10010) | 28 | 6.32 |
|  | Voltage-Gated Ca, Na (PNTHR10037) | 42 | 15.46 |
| PFAM | ABC Transporter (PF00005) | 23 | 4.51 |
|  | RNA recognition motif (PF00067) | 35 | 5.29 |
|  | Protein kinase domain (PF00069) | 30 | 4.58 |
|  | Response regulator receiver (PF00072) | 30 | 5.07 |
|  | Helicase C (PF00271) | 202 | 119.64 |
|  | AMP-binding enzyme (PF00501) | 62 | 8.93 |
|  | Binding protein dependent (PF00528) | 5 | 4.12 |
|  | Major Facilitator Superfamily(PF07690) | 75 | 13.86 |
| Nextstrain | SARS-CoV-2 | 428 | 13731 |
|  | Mumps | 232 | 238 |

Table indicates the number of identified monophyletic subfamilies and time for analysis.

## Discussion

In this study, we demonstrated the ability of our phylogenetic clustering approach AutoPhy to automatically identify monophyletic subfamilies in a wide range of protein families with highly differential rates of evolution. AutoPhy rapidly partitioned even very large and complex phylogenies into subfamilies without the need for arbitrary *ad hoc* parameter values. We also showed that AutoPhy log ratio analysis identifies highly divergent clades within subfamilies that may have been subjected to strong selection. In addition to protein subfamily clustering, AutoPhy also identified phylogenetic clusters in rapidly evolving viruses (e.g., SARS-CoV-2), and is highly robust to highly differential rates of evolution both among and within phylogenies.

AutoPhy rapidly clusters even very large phylogenetic trees (i.e., trees with many sequences), though the speed of the clustering is highly dependent on the tree topology. The time it took to cluster protein families into monophyletic groups ranged from 3 sec for the Ski oncogene, PNTHR10005 (628 protein sequences), to 2 min for the helicaseC protein family, PF00271, (421 protein sequences; Table 2). While one might expect that the process would take longer with bigger phylogenies, the tree topology, specifically the ratio of branch lengths within and between phylogenetic clusters, played a much bigger role in determining analysis time. For example, AutoPhy required less time to cluster the 1090 sequences in the PNTHR10037 phylogeny (16 sec; Table 2) than to cluster the 421 sequences in the helicaseC phylogeny (2 min; Table 2). The phylogenetic trees that took the shortest amounts of time to analyze tended to have discrete clusters separated by long branch lengths from other groups. The Ski oncogene tree that took only 3 sec to analyze contained many tightly bound subtrees with larger branch lengths leading to the cluster (suppl Fig. 1), whereas the smaller helicaseC phylogeny had much sorter subtending branches (suppl Fig 2). This pattern also held for the analyses of the virus phylogenies. The number of sequences in the SARS-CoV-2 phylogeny was less than twice the size of the Mumps phylogeny, but the analysis of the SARS-CoV-2 phylogeny took 57 times longer. A comparison of the two phylogenies reveals that there were many much more clearly, visually distinct, subtrees in the mumps phylogeny than in the SARS-CoV-2 phylogeny (suppl Figs 1 and 2).

A run time analysis of AutoPhy’s algorithmic workflow found that for most of the phylogenies the PDM creation took 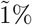 of the total time, while the UMAP took 23-28% of total time, GMMEM analysis required between 19-25% of the total time, and the MCIM required between 25-40% of the total run time. However, in the case of the trees with long run times (e.g., the SARS-CoV-2, Mumps and helicaseC phylogenies), 95% of the total run time was spent on the MCIM step. In other words, if the clusters from UMAP-GMMEM successfully identified monophyletic groups based on the PDM, very little time was spent on the MCIM step. The UMAP-GMMEM step appeared to have the most difficulty with polyphyletic sequences that were closely related and had long intra- but short inter-distances. For example, the UMAP-GMMEM analysis of the BSH tree clustered two groups as polyphyletic groups when related back to the tree, groups 1 and 8 (Fig 7A). In both cases, the polyphyletic clustering was due to a singleton sequence with a short internode distance (groups 1.1 and 8.1, Fig 7B). Similar patterns can be observed with the analysis of the OMP proteins (e.g., groups 7 and 8; Fig 4B, 6). The UMAP-GMMEM also had a particularly difficult task with polyphyletic clustering with the Acyltransferase (AT) phylogenetic tree. The phylogenetic clades within this tree tended to have long intranode branches within clades and separated by short internode branches between clades (Fig. 7C, D). In addition to difficulties resolving short internodes, much of the polyphyletic UMAP-GMMEM clustering was due to the inclusion of small groups of sequences or singletons. This suggests that greater sampling of these subfamilies would have improved the ability to resolve these clades.

These result support the idea that tree topology played a major role in determining overall run time. The UMAP method is a dimensional reduction tool that is especially good at optimizing low-dimensional graphs, ensuring that local structure is preserved [33]. Thus, the UMAP-GMMEM approach should be best at resolving monophyletic clusters within trees containing many discrete clades separated by long branches. For such trees, not much time would be spent resolving polyphyletic groupings with MCIM. Given these results, we suggest that improving the speed and efficiency of the MCIM algorithm, (e.g., by converting from Python to C), followed by speed improvements to UMAP-GMMEM, would provide the greatest overall run time decrease for AutoPhy, especially with difficult to resolve phylogenies like the SARS-CoV-2.

A detailed inspection of the clustering results of the well-studied and annotated protein families found that the AutoPhy clustering identified most of the functional subfamily annotations within the same monophyletic group, though in many cases these subfamilies included other closely related sequences. For example, AutoPhy placed the TOLCs efflux pumps that transfer small molecules such as antibiotics across bacterial outer membrane of bacteria [34] within a clade (group 8.0, Fig. 6) that also included the OMP5 and BEPC proteins, which are also known for their antibiotic resistance activity [35]. Other clusters included the 8 multidrug resistance outer membrane proteins MdtQ/P (group 0.0), and the 8 nucleoside-specific channel porins TSX (group 3.0). In other cases, the annotated sequences did not have enough sequences to form a discrete clade, such as the two OMPK sequences which are receptors for a broad host-range vibriophage KVP40 [36]. The OMPK grouped into a larger monophyletic group with OMPX, a small (18 kDa) outer-membrane protein. There were also a few cases in which AutoPhy split sequences with the same annotation into different clusters, such as the OMPA sequences, three (group 0.2) of which did not cluster with the larger OMPA group (group 0.0). A visual examination of the phylogeny shows that these three OMPAs were significantly evolutionarily divergent from the others, suggesting they may have evolved unique functional roles.

In other cases, AutoPhy identified interesting subfamily clusters within protein families which may indicate novel functional roles. For example, all the BSH sequences were annotated as choloylglycine hydrolases (Table 1). However, a visual inspection of the BSH tree clearly indicates the BSH phylogeny includes several distinct monophyletic groups (Fig. 7B) and AutoPhy split the sequences into probable subfamilies likely differing in kinetic characteristics or substrate activation. In the acyltransferases, most sequences with the same annotations (e.g., ceramide synthases (CERS)) were clustered by AutoPhy (group 0.0, Fig. 7B). One exception was the membrane-bound palmitoyltransferases (ZDH) which were split into two groups of only ZDH annotated sequences. The bigger AutoPhy-identified cluster of ZDH (group 11) are Golgi and endoplasmic reticulum bound and provide a variety of palmitoyl protein substrates [37]. The other ZDH group identified by AutoPhy (group 4.2) included ZDH14, which may regulate G-protein-couple receptor signaling (PubMed:27481942), ZDH11, a palmitoyltransferase involved in immune response, and ZH11B, a probable membrane-bound palmitoyltransferase involved in cell signaling in an immune response. The fact that separate ZDH clusters detected by AutoPhy contained distinct groups of functions indicates that AutoPhy can identify novel functional roles in previously annotated sequences.

Finally, in the AutoPhy analysis of the nuclear receptors from UniProt, proteins with the same annotations (e.g., BMP, PIAS, and PARQ) all clustered together with additional, closely related sequences (Fig. 7E,F). For example, the sequence with UniProt accession number O51E2 grouped inside a clade of PARQ sequences (group 3). Interestingly, O51E2 is likely functionally related to the PAQR proteins. Both are membrane receptors, but the PARQ proteins bind steroid receptors [38], while the O51E2 substrate is unknown. Based on these results, receptor molecules for PARQ and O51E2 are most likely very similar. We hypothesize that the O51E2 substrate is an aromatic lipid with a structure similar to steroid hormones.

In addition to the automated monophyletic clustering, AutoPhy includes downstream analyses for further investigation of these clusters such as a log ratio approach for analyzing the relative evolutionary divergence rates among clusters. The relative evolutionary divergence of identified AutoPhy clusters revealed by the log ratio analysis indicates AutoPhy determined monophyletic groups under potential purifying selection at the amino acid level. For example, group 9.1 in the OMP tree (Fig. 8B), groups 6 in the BSH tree (Fig. 8C), and group 4.2 in the AT tree (Fig. 8D) have the lowest within group divergences compared to the rest of the subfamilies withing the tree. The log ratio scores also reveal sharp differences in the evolutionary rates among the various protein families.

While AutoPhy overcomes many limitations of previous approaches, this is room for improvement in several areas. Autophy has a tendency over split clades, particularly when there are long branches such as with the Acetyltransferases (Fig. 7D). AutoPhy analysis also produced many cluster groups containing only one or two sequences, but this would likely be resolved with greater sequence sampling. In addition to speed improvements noted earlier, other improvements to AutoPhy might include optimizing the number of UMAP dimensions for GMMEM clustering and including additional analysis to identify functional singletons in groups. Likewise, gaps could be added to the transition matrix to improve its estimated gap probability at sites. Lastly, the code is written in R and Python notebooks. This is excellent for demonstration and reproducibility but for creating databases or large-scale implementation needs to be configured as a python package as a command-line tool.

In addition to novel protein subfamily function identification and viral strain analysis, there are numerous other potential applications for AutoPhy. First, AutoPhy could be applied to metagenomic datasets to phylogenetically bin sequences by subfamily in a feature count table. This would allow for more refined metagenomic questions to be asked involving differentially present subfamilies. For example, a comparison of BSH abundance at the family level between conditions (e.g., healthy vs. dysbiotic gut microbiome samples) might not reveal any differences. However, in the same samples we might see a clear difference in the abundances of specific AutoPhy identified subfamily clusters within the BSHs. AutoPhy could also prove useful for automatically identifying different novel subfamilies present in the genomes of the same organisms, thereby identifying paralogs and functional misannotations. Finally, Autophy could also refine protein-matching. AutoPhy clusters could be used at the basis for subfamily-HMM models to refine protein matching to finer annotation.

## Supporting information

**S1 Fig. Ski oncogene tree AutoPhy analysis**.

**S2 Fig. Helicase C tree AutoPhy analysis**.

**S3 Fig. Mumps tree AutoPhy analysis**.

**S4 Fig. Covid tree AutoPhy analysis**.

## Acknowledgements

We would like to thank Nick Allsing for running our computers as the system admin and Rob Edwards for the computational power of his Anthill cluster.

