## Supplementary figures and images for "AutoPhy: Automated phylogenetic identification of novel protein subfamilies"

### supp_fig_1.tiff

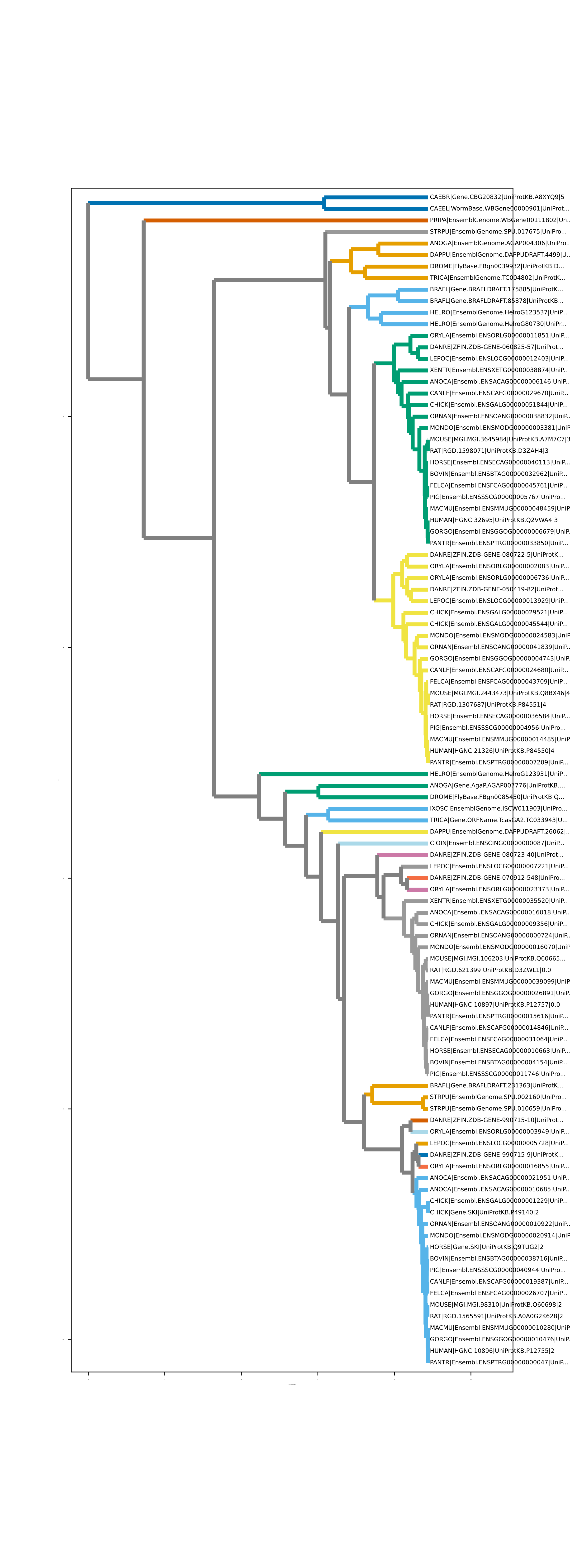

### supp_fig_2.tiff

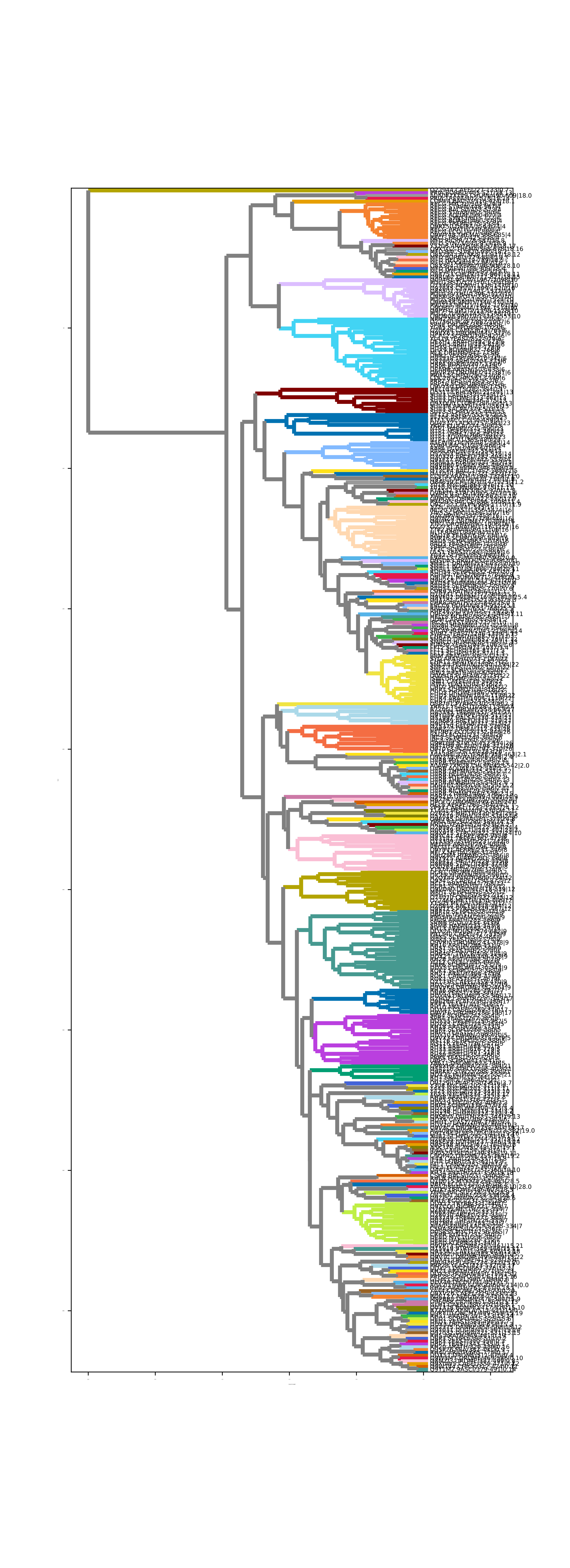

### supp_fig_3.tiff

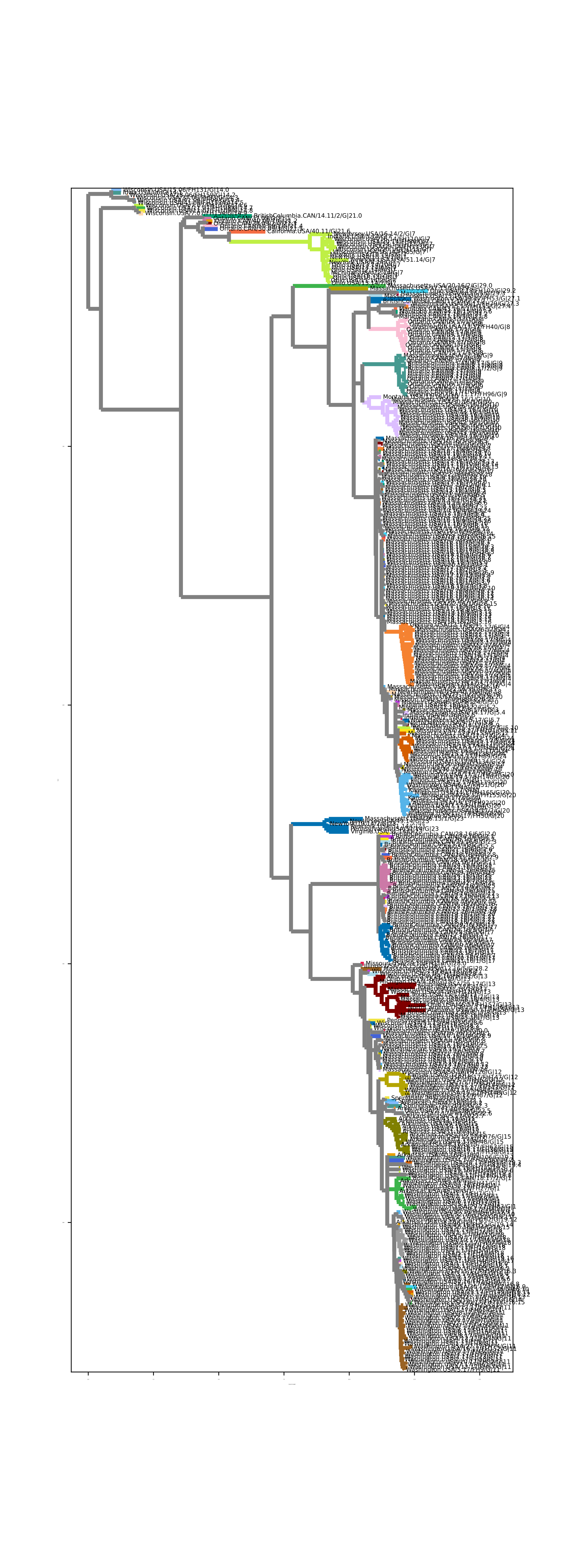

### supp_fig_4.tiff

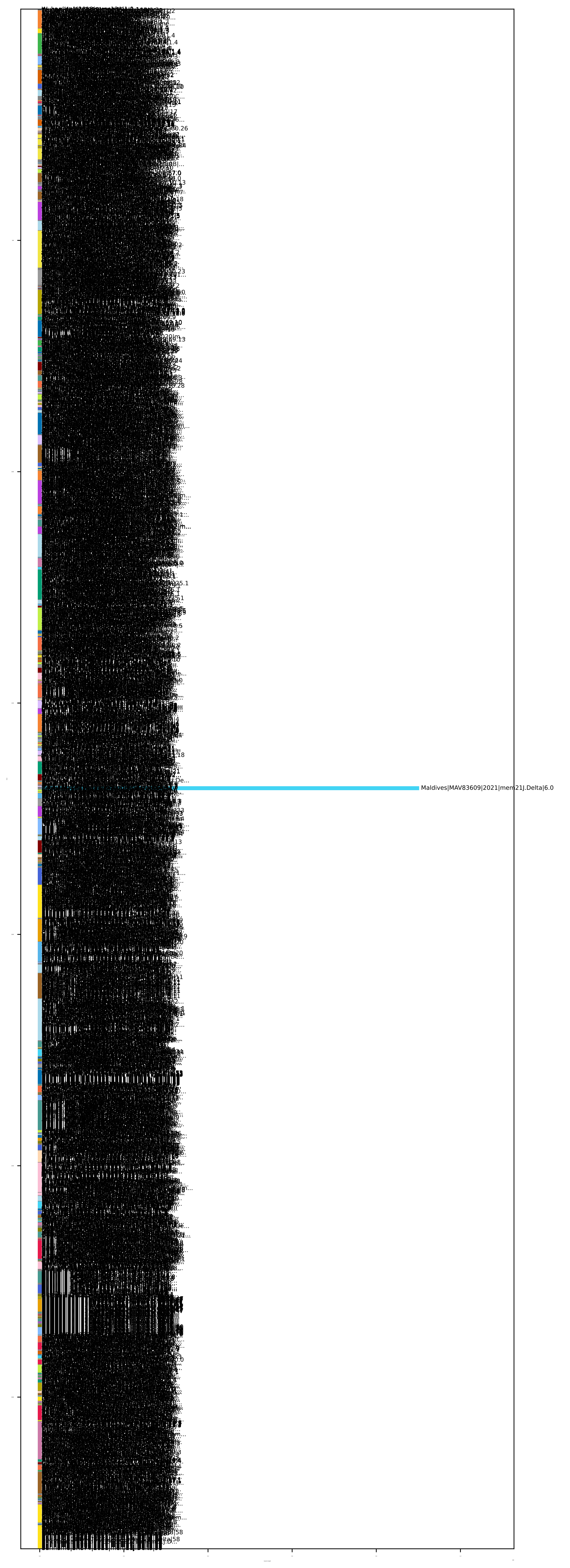
